# FN1 is produced by cancer-associated fibroblasts and carries no immune-exclusion signal beyond the collagen program: a two-cohort single-cell re-examination in non-small cell lung cancer

**DOI:** 10.64898/2026.11.23.768752

**Authors:** Zhibin Guo, Wenwu Bai, Xinchao Cao, Haiyan Jia, Liuqing Xing

## Abstract

Fibronectin 1 (FN1) has been proposed as a macrophage-derived driver of T-cell exclusion and AP-1– mediated resistance to checkpoint blockade in lung cancer. The human evidence was never tested against the two confounders most likely to generate them: collagen/extracellular-matrix (ECM) abundance and dissociation-induced immediate-early gene (IEG) expression. We reanalyzed two public non-small cell lung cancer (NSCLC) cohorts, GSE148071 (42 treatment-naive tumors) and GSE207422 (15 specimens; 92,330 cells), with 24 pre-treatment bulk transcriptomes. Malignant epithelium was called by compartment-level inferCNV; FN1 source was attributed within macrophage and fibroblast compartments and by ligand–receptor scoring; exclusion associations were tested by partial Spearman correlation with permutation testing, controlling for a 22-gene collagen/ECM signature; AP-1 regulon activity was quantified before and after masking the 27-gene IEG panel, benchmarked against a dissociation-stress score. FN1 was produced predominantly by cancer-associated fibroblasts (CAFs): the FN1-positive fraction was higher in CAFs than macrophages in both cohorts (paired Wilcoxon P = 1.6 × 10⁻⁹ and P = 0.002), and CAF-to-epithelium communication exceeded macrophage-to-epithelium by about six-fold (median 0.32 vs 0.05, P = 1.5 × 10⁻⁵). An independent lung-adenocarcinoma cohort (GSE131907) replicated this cell of origin (FN1-positive fraction 0.88 in CAFs versus 0.44 in TAMs; 11 paired samples, P = 0.001). The bulk FN1–cytotoxic association (ρ = −0.53, P = 0.008) disappeared after collagen adjustment across three independent FN1 scoring schemes (partial ρ = +0.21 to −0.16, all P > 0.33) and was not reproduced at single-cell level. Sample-level AP-1 activity tracked dissociation stress (ρ = 0.88, P = 2.3 × 10⁻⁹); IEG masking shrank histology-associated effect sizes by 40–64%, yet masked activity remained stress-coupled (ρ = 0.82). FN1 is therefore a CAF product whose apparent link to immune exclusion is absorbed by the collagen program, and AP-1 readouts from dissociated tissue are stress-dominated; stress scores should accompany AP-1-based resistance claims.

**Author Summary:** Lung tumours are not just cancer cells. They also contain fibroblasts, which build a fibrous scaffold, macrophages, which normally engulf microbes and debris, and T cells, which can kill tumour cells. A protein called fibronectin, encoded by the FN1 gene, has been reported to be made by macrophages inside lung tumours and to keep T cells out of the tumour, which would make it an attractive drug target. We asked whether the available human data actually support that account. Using two published collections of lung-cancer samples, one profiled cell by cell and the other profiled as bulk tissue, we found that fibronectin is produced mainly by fibroblasts rather than by macrophages. In bulk tissue, tumours containing more fibronectin did have fewer cytotoxic T cells, but this relationship disappeared once we accounted for how much collagen and other matrix the same tumours contained; fibronectin was largely travelling with the fibrous scaffold. We also found that a widely used readout of tumour aggressiveness, the activity of the AP-1 genes, is inflated by the way samples are handled in the laboratory. Our results do not overturn the biology of fibrosis in cancer, but they argue that fibronectin and AP-1 measurements in human tumours should not be treated as independent drug targets or biomarkers until collagen content and sample-handling stress are taken into account.

## Introduction

Fibronectin 1 (FN1), an extracellular-matrix glycoprotein, has attracted attention as a candidate mediator of immunotherapy resistance in solid tumors. A recent study combined macrophage-specific Fn1 knockout in mouse models with single-cell transcriptomics and multiplex imaging of human non-small cell lung cancer (NSCLC) specimens, and concluded that tumor-associated macrophages (TAMs) are a major FN1 source that promotes matrix stiffening, immunosuppressive macrophage polarization and impaired T-cell infiltration [1]. Single-cell analyses of anti-PD-1-treated lung cancer have in parallel nominated JUN and related AP-1 factors as response biomarkers [2], positioning the AP-1 family as the downstream effector that would link matrix-derived integrin signals to checkpoint-resistance phenotypes.

Two obstacles make this narrative difficult to evaluate in human tissue. The first is attribution. Bulk RNA sequencing cannot resolve which cell type produces a ligand, and FN1 is co-expressed with collagen and other matrix genes in stromal compartments. An association between tumor-level FN1 and immune exclusion may therefore reflect nothing more than the stromal content of the sample; without an explicit adjustment for collagen-program abundance, the specific contribution of FN1 cannot be separated from that of the extracellular matrix in general. The human evidence for the macrophage-source model has not, to our knowledge, been tested against this compositional confounder, nor against the fibroblast compartment as a quantitative alternative source.

The second obstacle concerns the readout. JUN, FOS and related AP-1 factors are immediate-early genes (IEGs) induced within minutes of tissue dissociation [3–5]. In single-cell preparations, AP-1 activity measured in tumor epithelium is therefore a composite of genuine integrin-driven signaling and a processing artifact whose magnitude varies from sample to sample. Because the proposed FN1–AP-1 axis uses AP-1 activity as its downstream endpoint, any test of the axis must first quantify and remove the dissociation component.

Here we address both obstacles using two complementary public cohorts: GSE148071, a treatment-naive NSCLC single-cell atlas containing both lung adenocarcinoma (LUAD, n = 18) and squamous cell carcinoma (LUSC, n = 24 as called here) [6], and GSE207422, which provides single-cell profiles of 15 resected or biopsied specimens with pathological-response annotation after neoadjuvant PD-1 blockade, plus 24 pre-treatment bulk transcriptomes [7]. We quantify the cellular source of FN1, test whether its association with T-cell exclusion survives adjustment for the collagen program, and measure how much of the reported AP-1 signal is attributable to dissociation stress.

## Methods

### Data sources and preprocessing

Count matrices were obtained from GEO (GSE148071; GSE207422). GSE148071 contains raw UMI counts (genes × cells, tab-separated) for 42 advanced NSCLC tumors [6]. GSE207422 contains a raw UMI matrix (24,292 genes, 92,330 cells, BD Rhapsody whole-transcriptome amplification) for 15 specimens (12 resections after neoadjuvant anti-PD-1 therapy and 3 pre-treatment biopsies) and a log2TPM bulk matrix for 24 pre-treatment tumors from an independent treatment-naive set [7]. Clinical annotations (treatment group, histology, pathological response, sampling time) were parsed from the series matrix files. A third public cohort, GSE131907 [11], was used as an independent replication set for the FN1 cell-of-origin analysis only: its raw UMI matrix (29,634 genes, 208,506 cells from 58 lung, lymph-node and brain-metastasis specimens) and cell-level annotations were downloaded from GEO, and the published cell-type labels (Fibroblasts, Myeloid cells) were used without re-annotation. Per-sample cell and gene counts before and after quality control are provided in Tables S1 and S2.

### Cell-type annotation and quality control

Cells with fewer than 200 detected genes or a mitochondrial fraction above 0.25 were excluded (12,537 cells, 13.9%, in GSE148071). Remaining cells were assigned to compartments (epithelial, myeloid, lymphoid/immune, fibroblast, endothelial, unassigned) by mean marker-score comparisons using canonical markers (epithelial: EPCAM, KRT8/18/19, KRT5/17; myeloid: LYZ, CD68, CD14, CSF1R; lymphoid: CD3D, CD3E, NKG7, MS4A1; fibroblast: COL1A1, COL1A2, DCN, LUM, POSTN; endothelial: PECAM1, VWF, CLDN5). The published GSE148071 series reports 18 LUAD, 22 LUSC and 2 NSCLC-not-otherwise-specified samples [6]; the two NOS samples were reassigned to LUSC by epithelial marker inference (squamous: KRT5, KRT6A, KRT17, TP63, DSC3; adenocarcinoma: NAPSA, SFTPC, SLC34A2, MUC1), giving the 18 LUAD / 24 LUSC split used throughout. Inferred assignments with ambiguous marker scores (3 samples) were retained in compartment-level analyses and flagged in Table S1.

### Malignant-cell calling

Malignant epithelial cells were identified with a purpose-built inferCNV implementation (NumPy/SciPy): expression was normalized to counts-per-10,000 followed by log1p transformation, gene orders were taken from NCBI Gene annotation, smoothed residuals were computed in 100-gene sliding windows against a single-cell-type reference (immune cells by preference), and per-cell CNV scores were aggregated to a compartment-level consensus chromosome-arm profile. A sample was called malignant when at least four chromosomes exceeded max(4σ of the reference null, 0.015); reference pools were required to contain at least 30 cells. Single-type references were used throughout because mixed reference pools produce chimeric profiles and near-universal false-positive calls. In GSE148071, 38 of 42 samples were CNV-evaluable; 26 were called malignant. In GSE207422, 10 of 15 samples were called malignant; the single pathological-complete-responder was called CNV-negative, supporting specificity (Fig. S1). Two sample sets are used downstream: the full set of 26 CNV-positive GSE148071 samples entered the regulon analysis, and a 23-sample subset yielding at least 50 malignant epithelial cells (LUAD 9, LUSC 14) served as a prespecified sensitivity set (Fig. 1e).

**Figure 1.**
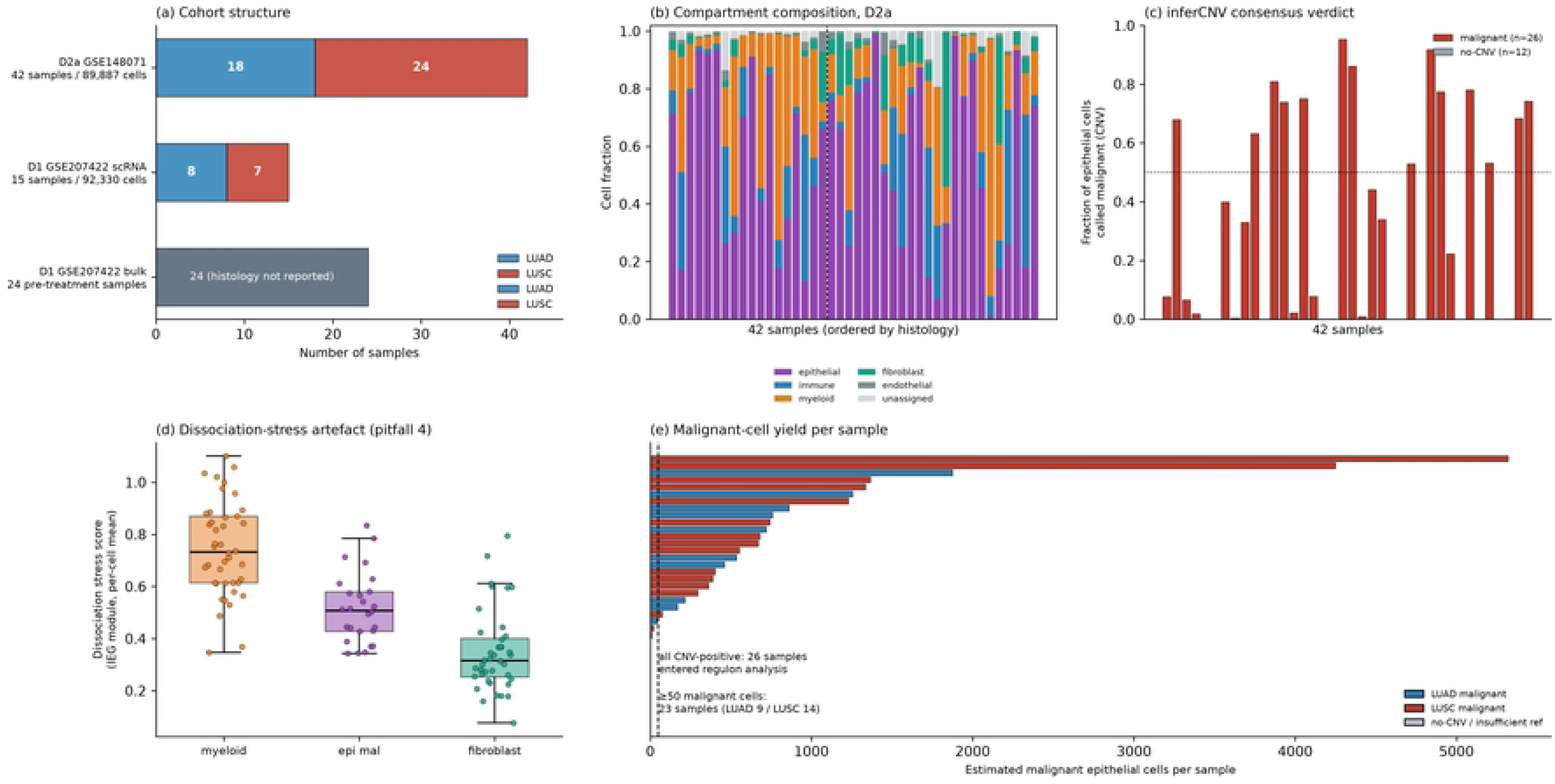
Two-cohort design, cell atlas, malignant-cell calling and the dissociation-stress audit. (a) Cohort structure across GSE148071 (42 treatment-naive tumors) and GSE207422 (15 single-cell specimens and 24 pre-treatment bulk samples). (b) Compartment composition of the 42 GSE148071 samples, ordered by histology. (c) Compartment-level inferCNV consensus verdicts with the fraction of epithelial cells called malignant per sample. (d) Dissociation-stress scores by compartment, ordered as in the text (myeloid, malignant epithelium, fibroblast). (e) Estimated malignant-cell yield per sample; 26 CNV-positive samples entered regulon analysis, of which 23 (LUAD 9 / LUSC 14) yielded at least 50 malignant cells.

### Dissociation-stress scoring and AP-1 quantification

A per-cell stress score was computed as the mean normalized expression of a 27-gene dissociation-stress/IEG panel (FOS, FOSB, FOSL1, FOSL2, JUN, JUNB, JUND, EGR1, EGR2, EGR3, ATF3, HSPA1A, HSPA1B, HSPA6, HSPB1, DNAJB1, HSPH1, HIST1H1C, BTG2, CSRNP1, DUSP1, KLF4, NR4A1, NR4A2, NR4A3, RHOB, ZFP36) [3–5], aggregated per sample and compartment as the mean of per-cell scores within each sample–compartment set (malignant-epithelium scores required at least 20 cells). Regulatory activity was inferred with a SCENIC-style pipeline implemented in NumPy/SciPy [8]: within each sample, Pearson correlations between transcription factors (curated list) and all expressed genes were Fisher-z-transformed and sample-size-weighted; edges with weighted ρ ≥ 0.10 (top 100 targets per TF, minimum 20 targets) defined regulons; regulon activity was scored with an AUCell-like top-5% recovery metric. Cells entering this pipeline additionally required more than 500 total counts. Two versions were computed for every regulon: a full version and an IEG-masked version in which the same 27 panel genes were excluded from target sets. A third, hard-masked version removed the same 27 genes from the gene universe used to build the expression window (29,527 → 29,500 genes), so that panel genes could not occupy window slots at all, and recalculated the window size accordingly (top 5%: 1,476 → 1,475 genes per cell). The FOSB regulon lost its scoring eligibility after masking (11 of its 30 targets are IEG-panel members, leaving fewer than the 20-target minimum) and is therefore reported as full-version only. This yielded 14 full and 13 masked regulons. Analyses were run on malignant epithelial cells from the 26 CNV-positive GSE148071 samples (43,324 cells; LUAD 11 / LUSC 15 samples contributed cells). Per-sample regulon statistics and activities are provided in Tables S3 and S6 and Fig. S3.

### Ligand–receptor communication

Fourteen curated ligand–receptor pairs centered on the FN1/integrin axis (FN1 with CD44, ITGA5-ITGB1, ITGAV-ITGB1, ITGA9, ITGA4; HGF–MET; TGFB1–TGFBR1/TGFBR2; SPP1–CD44 and SPP1–ITGAV-ITGB1; CCL2–CCR2; CXCL12–CXCR4; COL1A1–ITGA2-ITGB1; TNC–ITGAV-ITGB1; VIM–CD44) were scored per sample as the product of mean ligand expression in the ligand compartment (TAM or CAF) and mean receptor expression in malignant epithelial cells; samples contributing fewer than 30 cells to a compartment were not scored for that compartment. A sample-level FN1-axis score was defined as the mean of the five FN1-containing pairs. CAF versus TAM contributions were compared by paired Wilcoxon signed-rank tests on samples with both compartments. Histology contrasts used Mann–Whitney U tests with Bonferroni correction across the 14 pairs within each ligand compartment. Sample-level scores are provided in Fig. S2 and Tables S4 and S7.

### Bulk-tissue signatures and attribution analysis

For the 24 pre-treatment bulk samples, signature scores were computed as the mean log2TPM of: an FN1-axis signature (FN1 together with six ECM-remodeling genes co-expressed in the stromal program — COL1A1, COL11A1, SPP1, MMP2, LOX and TNC); FN1 alone; a collagen-free FN1-receptor program (the mean of the FN1 receptors detectable in bulk — ITGA5, ITGB1, ITGAV, ITGB5, ITGA9, ITGA4, CD44, SDC1, SDC4 and LRP1 — excluding FN1 itself and all collagen/ECM genes); a 22-gene collagen/ECM signature (excluding FN1); an epithelial–mesenchymal transition signature [9]; a proliferation signature; and a cytotoxic T-cell signature (CD8A, GZMB, PRF1, IFNG, NKG7, GZMA), following cytolytic-activity marker conventions [10]. Associations with the cytotoxic signature were tested by partial Spearman correlation: variables were rank-transformed, residuals were taken against the covariate set by ordinary least squares, and permutation P values (5,000 draws) were computed for the residual correlation. The same framework was applied to single-cell sample-level variables (FN1-positive fractions, stress scores, compartment scores) in both cohorts, with raw, ECM-adjusted, and ECM-plus-stress models; the single-cell cytotoxic signature used for GSE207422 attribution analyses comprised GZMA, GZMB, GZMK, GZMH, PRF1, GNLY, NKG7, IFNG, KLRD1, CST7 and FGFBP2. Compartment-level summaries entering these models were subject to sample-level size minima — at least 20 fibroblasts for CAF-based summaries and 50 myeloid cells for TAM-based summaries — independent of the 30-cell minimum applied to ligand– receptor scoring above. Because GSE148071 contains very few immune cells (5,439 cells; median 66 per sample), immune-exclusion analyses in that cohort were restricted to density diagnostics and are reported as such.

### Statistical analysis

Non-parametric tests were used throughout (Mann–Whitney U, Wilcoxon signed-rank, Spearman correlation) with two-sided P values. Multiple testing was handled by Bonferroni correction within families of hypotheses, as stated. All analyses were implemented in Python 3 (NumPy, pandas, SciPy, Matplotlib); analysis scripts and per-sample result tables are archived at Zenodo (DOI 10.5281/zenodo.24013282). No sample was excluded other than by the prespecified quality-control and eligibility rules above.

## Results

### Two cohorts with an explicit stress audit

The treatment-naive cohort (GSE148071) contributed 89,887 cells across 42 tumors (77,350 after quality control) with rich epithelial and stromal content (Fig. 1a,b); Table 1 summarizes the three datasets and the role each plays in the analysis. Compartment-level inferCNV consensus calling identified malignant epithelium in 26 of 38 evaluable samples; 23 samples (LUAD 9, LUSC 14) yielded at least 50 malignant cells (Fig. 1c,e; Fig. S1). The treated cohort (GSE207422) contributed 92,330 cells, dominated by immune (54,298) and myeloid (22,774) populations with 13,291 epithelial cells, of which malignant epithelium was detected in 10 of 15 samples (Fig. 1c). A built-in specificity check passed: the single pathological-complete-responder sample was called CNV-negative, consistent with effective clearance of malignant cells by therapy.

**Table 1.** Cohort and sample summary. (See Tables S1 and S2 for per-sample values.)

| Dataset | Platform | Samples | Cells | LUAD/LUSC | Role in analysis |
|---|---|---|---|---|---|
| GSE148071 | Raw UMI (tab-separated) | 42 tumors | 89,887 (77,350 post-QC) | 18 / 24 (2 NOS reassigned to LUSC by marker inference) | FN1 source, histology contrasts, regulons |
| GSE207422 scRNA | BD Rhapsody WTA, raw UMI | 15 specimens (12 post-treatment, 3 pre-treatment) | 92,330 | 8 / 7 | Replication, response contrasts, immune-side analyses |
| GSE207422 bulk | log2TPM | 24 pre-treatment tumors | — | not reported | Immune-exclusion and attribution analyses |

Dissociation stress was non-uniform across compartments: myeloid cells showed the highest IEG module scores (median across samples 0.732), malignant epithelium intermediate (0.508) and fibroblasts lowest (0.316) (Fig. 1d; scores are per-sample means of per-cell values). Any cross-sample comparison of AP-1 activity therefore requires an explicit correction, which we implement below.

### FN1 is produced predominantly by cancer-associated fibroblasts

Within compartments, the fraction of cells expressing FN1 was markedly higher in fibroblasts than in macrophages in both cohorts. In GSE148071, the median FN1-positive fraction was 0.80 in LUAD and 0.63 in LUSC CAFs versus 0.31 and 0.09 in TAMs (paired Wilcoxon P = 1.6 × 10⁻⁹ across 40 paired samples); in GSE207422 the same direction held (CAF median 0.85 versus TAM 0.40, P = 0.002 across 11 paired samples) (Fig. 2a,b). Ligand–receptor scoring gave the same answer at the communication level: sample-level CAF-to-epithelium FN1-axis scores (median 0.32) exceeded TAM-to-epithelium scores (median 0.05) approximately six-fold (paired Wilcoxon P = 1.5 × 10⁻⁵, 17 samples with both compartments) (Fig. 2c,d); in the six GSE207422 samples with both compartments represented, the same CAF-over-TAM contrast held (P = 0.031; Fig. 2e). The FN1-positive fraction of TAMs was itself histology-dependent, being approximately 3.6-fold higher in LUAD than in LUSC (0.304 versus 0.084, P = 0.0004; Fig. 3a), a pattern consistent with the possibility that macrophage-derived FN1 reports in adenocarcinoma-rich series partially reflect cohort composition. The cell-of-origin result was further replicated in an independent public LUAD cohort (GSE131907; 208,506 cells from 58 specimens) using the deposited raw counts and published cell-type labels: in the 11 tumour samples with at least 20 cells in both compartments, the median FN1-positive fraction was 0.88 in fibroblasts versus 0.44 in myeloid cells (paired Wilcoxon P = 0.000977; approximately twofold), and the same direction held across all 24 samples with sufficient cells (0.83 versus 0.46, P = 5.96 × 10⁻⁷; S9 Table). Normal-lung samples showed no consistent compartment difference, consistent with a tumour-associated stromal source.

**Figure 2.**
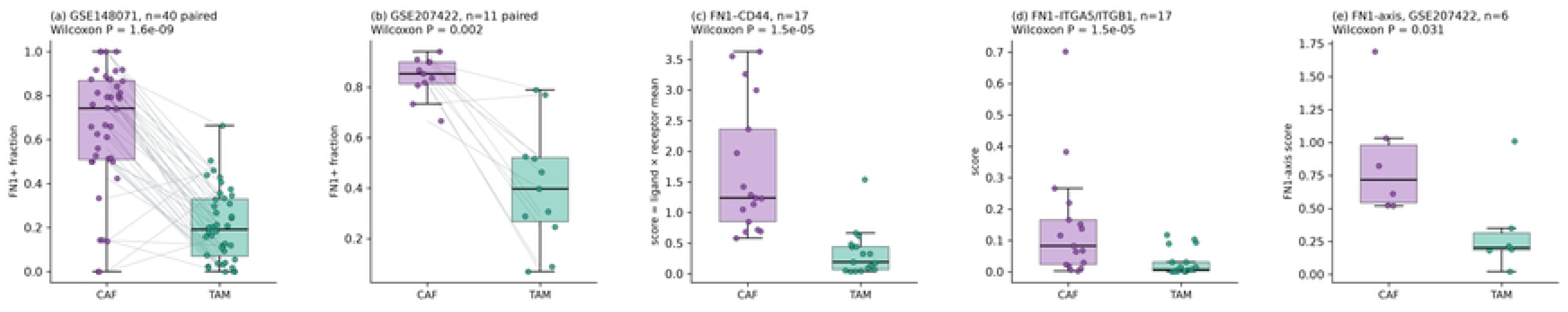
FN1 is produced by cancer-associated fibroblasts. (a) FN1-positive fraction in paired CAF and TAM compartments per sample, GSE148071 (paired Wilcoxon test). (b) FN1-positive fraction in paired CAF and TAM compartments per sample, GSE207422. (c, d) Sample-level FN1–CD44 and FN1– ITGA5/ITGB1 communication scores with TAM versus CAF as the ligand compartment. (e) The same ligand–receptor contrast in GSE207422 (n = 6 samples with both compartments represented; paired Wilcoxon P = 0.031).

**Figure 3.**
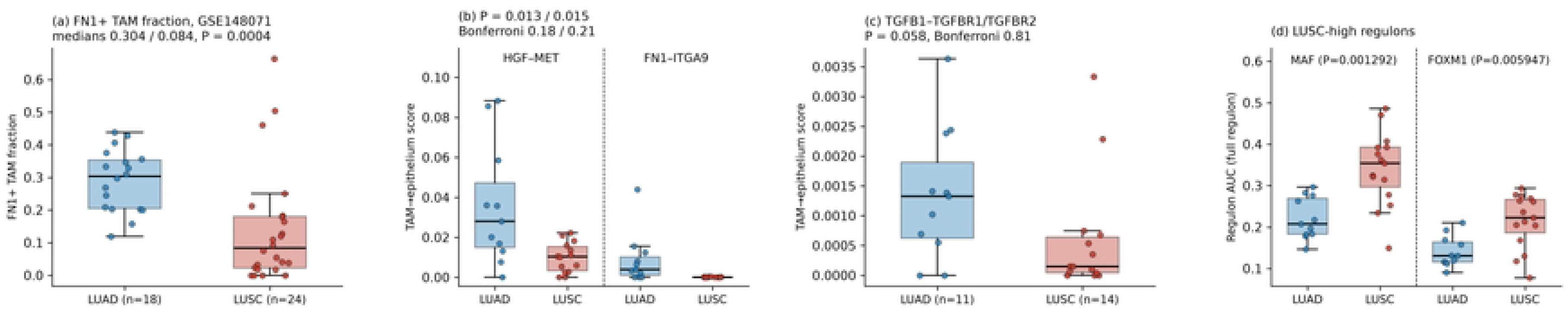
Histology contrasts in the FN1 axis and downstream regulons. (a) FN1-positive TAM fraction in LUAD versus LUSC. (b, c) HGF–MET, FN1–ITGA9 and TGFB1–TGFBR1/2 communication scores by histology (TAM-to-epithelium; none of the 14 pairs survives Bonferroni correction). (d) MAF and FOXM1 regulon activity by histology.

### The FN1–exclusion association is absorbed by the collagen program

In pre-treatment bulk tumors, FN1-axis scores correlated negatively with the cytotoxic T-cell signature (ρ = −0.53, P = 0.008), as did FN1 alone (ρ = −0.47, P = 0.020) and a collagen-free program built from the FN1 receptors only, excluding FN1 itself (ρ = −0.55, P = 0.005). After adjustment for the 22-gene collagen/ECM signature, all three associations disappeared (partial ρ = +0.12, +0.21 and −0.16 respectively; all P > 0.33; Fig. 4a,b; Table S5). Because the collagen-free signature shares no genes with the covariate, this collapse does not reflect a circular adjustment.

**Figure 4.**
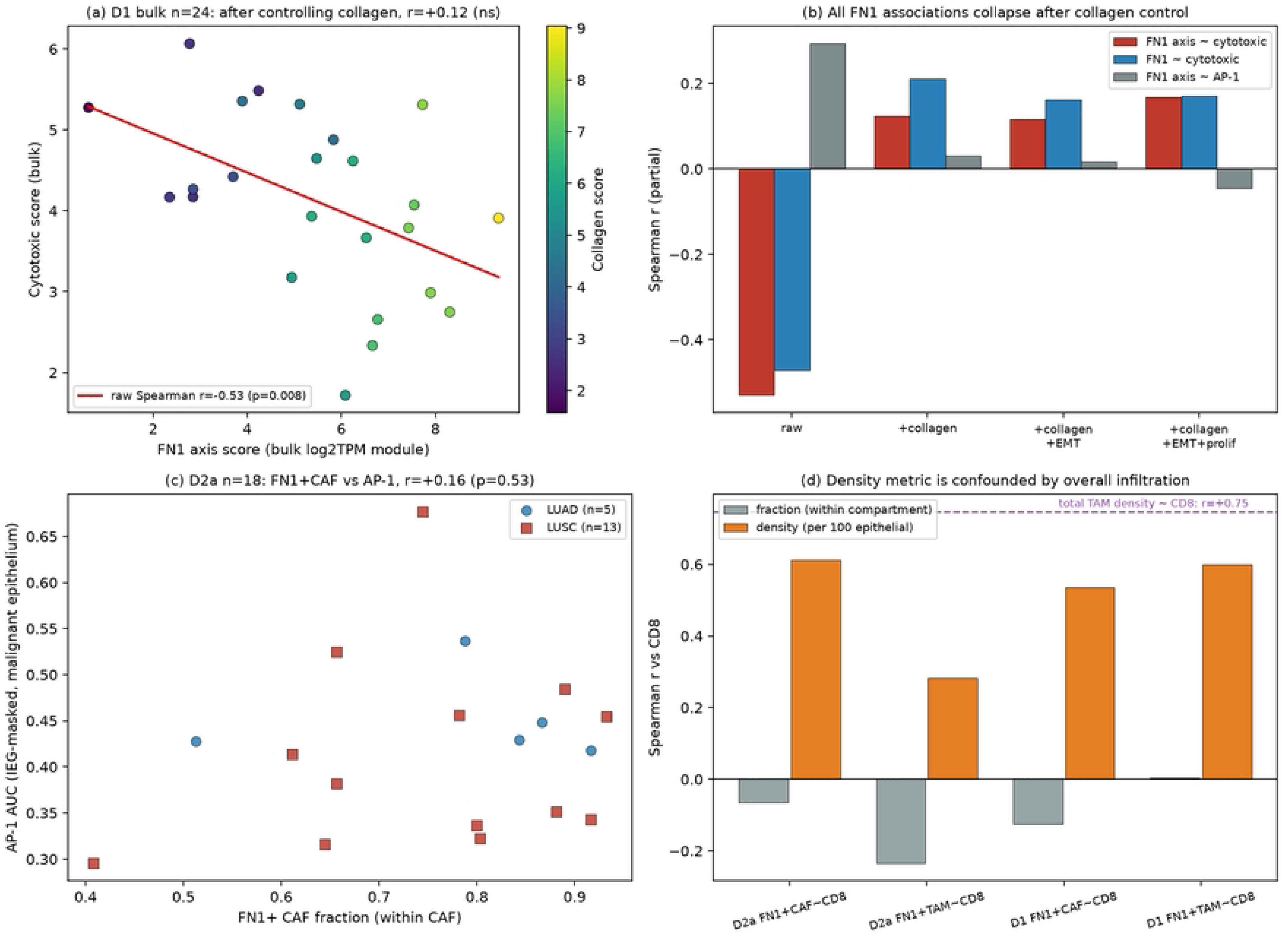
Attribution analysis: FN1 versus the collagen program. (a) Bulk-tissue association between the FN1-axis signature and the cytotoxic signature in 24 pre-treatment tumors, with the partial correlation after collagen adjustment. (b) Raw and partially adjusted Spearman correlations for the three FN1 signatures against cytotoxic and AP-1 readouts. (c) FN1-positive CAF fraction versus IEG-masked AP-1 activity in GSE148071 malignant epithelium (n = 18 of the 26 CNV-positive samples: samples with at least 20 CAFs and an estimable FN1⁺ fraction; ρ = +0.16, P = 0.53). (d) Fraction-versus density-based correlations with CD8 abundance, illustrating confounding by total infiltration.

Single-cell data did not rescue the association. In GSE148071, immune cells were too few for inference (median 66 per sample). In GSE207422, the direction reversed: the FN1-positive fraction of TAMs correlated positively with within-sample cytotoxic activity (ρ = +0.71, P = 0.003, n = 15), and the correlation remained positive after ECM adjustment (partial ρ = +0.80, P = 0.006). Density-based measures were dominated by overall infiltration: total TAM density correlated with CD8 density more strongly (ρ = +0.75) than FN1-positive TAM density did (ρ = +0.61) (Fig. 4c,d). The bulk-level negative association thus reflects the tissue-composition effect that stroma-rich tumors are immune-cold; once composition is removed, FN1-positive status tracks immune activation rather than exclusion. We conclude that FN1 behaves as a marker of collagen-high CAF programs and carries no immune-exclusion information beyond the matrix program in human NSCLC.

### AP-1 readouts are stress-dominated and require more than IEG masking

Sample-level AP-1 regulon activity in malignant epithelium tracked the per-sample dissociation-stress score closely (Spearman ρ = 0.88, P = 2.3 × 10⁻⁹, n = 26; Fig. 5d), confirming that measured AP-1 activity is largely a processing artifact. Masking the 27 stress-panel genes from regulon targets reduced histology-associated AP-1 effect sizes by 40–64% (JUN: log2FC 0.28 → 0.11; JUNB: 0.30 → 0.18; FOS: 0.29 → 0.17; ATF3: 0.28 → 0.10) (Fig. 5a,b; Table S3). After masking, AP-1 activity still did not differ between LUAD and LUSC (P = 0.64; the full-regulon contrast was likewise non-significant, P = 0.50; Fig. 5c).

**Figure 5.**
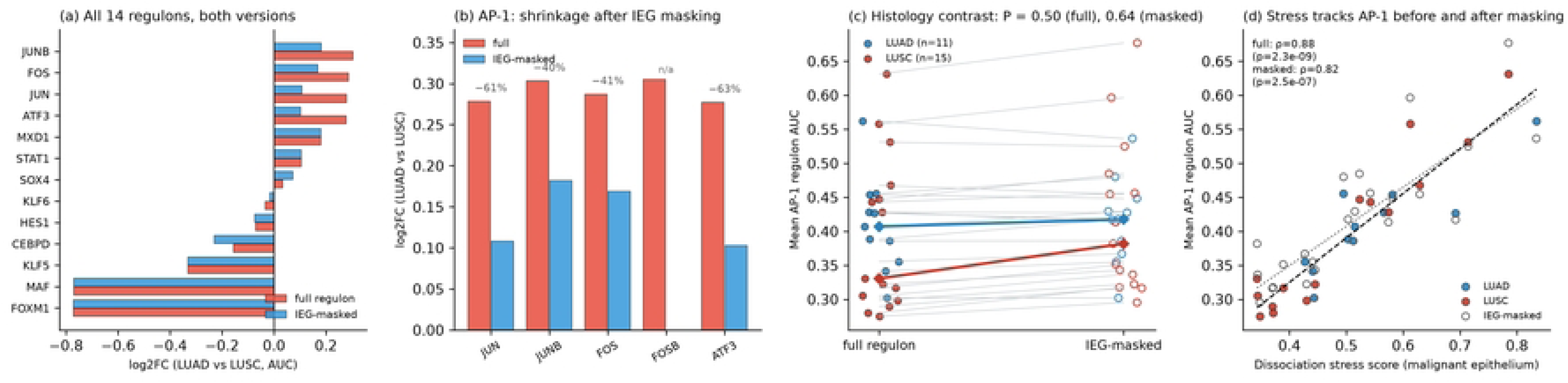
Immediate-early-gene masking shrinks the histology effect but does not remove stress coupling. (a) log2FC (LUAD versus LUSC) for all 14 regulons under full and IEG-masked versions. (b) AP-1 family effect sizes before and after masking, with percentage shrinkage; FOSB could not be scored after masking (n/a). (c) Mean AP-1 regulon AUC by histology under full and masked versions (P = 0.50 and P = 0.64). (d) Sample-level stress score versus AP-1 activity for full (ρ = 0.88) and masked (ρ = 0.82) versions, n = 26; a hard-masked version, in which the stress genes were removed from the ranking window, gave the same coefficient (Table S8).

Masking, however, is not a complete correction. IEG-masked AP-1 activity remained strongly coupled to the dissociation-stress score (ρ = 0.82, P = 2.5 × 10⁻⁷, n = 26; Fig. 5d), and the coupling was unchanged in the 23-sample sensitivity set restricted to samples with at least 50 malignant cells (ρ = 0.88 full and 0.82 masked). Masking removes the direct contribution of IEG transcripts to regulon target sets but cannot remove stress-driven AP-1 transcription-factor activation itself, which propagates through non-IEG targets. Purging the 27 panel genes from the ranking gene universe (29,527 → 29,500 genes), rather than only from the target sets, changed per-sample AP-1 activity by at most 0.0014 AUC units and left the stress coupling unchanged (ρ = 0.82, P = 2.5 × 10⁻⁷; Table S8), so the residual association is not an artifact of stress genes occupying window slots. Sample-level stress scores should therefore accompany any AP-1-based claim. Two non-AP-1 regulons showed histology differences among the 14: MAF was higher in LUSC and survived Bonferroni correction (P = 0.0013, adjusted P = 0.018; Fig. 3d), while FOXM1 showed the same direction but did not reach the corrected threshold (P = 0.006, adjusted P = 0.083; Fig. 3d). In the 23-sample sensitivity set the MAF contrast attenuated to adjusted P = 0.071, so this finding should be considered provisional. Prior single-cell reports of elevated epithelial AP-1 in therapy-resistant or squamous tumors [2] should be interpreted with the stress component in mind.

### Candidate communication axes

Among the 14 curated pairs scored with TAMs as the ligand compartment, three showed nominal or near-nominal elevation in LUAD: HGF–MET (P = 0.013), FN1–ITGA9 (P = 0.015) and TGFB1– TGFBR1/TGFBR2 (P = 0.058) (Fig. 3b,c; Table S4). None survived Bonferroni correction across the 14 pairs (adjusted P = 0.18, 0.21 and 0.81). The same was true with CAFs as the ligand compartment (smallest adjusted P = 0.081 for HGF–MET). Two further caveats apply. First, the ligand-side permutation test for FN1–ITGA9 indicated that FN1 is not enriched in the TAM compartment, consistent with the CAF origin established above. Second, these contrasts derive from a single cohort without independent replication, and their collinearity with stromal content was not exhaustively excluded. We therefore present them as hypothesis-generating only. Table 2 collects the key quantitative results and the statistic behind each claim.

**Table 2.** Key quantitative results.

| Question | Result | Statistics |
|---|---|---|
| FN1 source | CAF ≫ TAM (both cohorts) | FN1<sup>+</sup> fraction 0.63–0.85 vs 0.09–0.40; paired Wilcoxon P = 1.6 × 10<sup>−9</sup> (n = 40) and 0.002 (n = 11); FN1-axis communication ≈6-fold higher from CAFs (medians 0.32 vs 0.05; P = 1.5 × 10<sup>−5</sup>, n = 17) |
| FN1–exclusion (bulk) | Collapses after collagen adjustment | ρ = −0.53 → +0.12 (FN1-axis); −0.47 → +0.21 (FN1); −0.55 → −0.16 (collagen-free); all P > 0.33 |
| FN1–exclusion (single-cell) | Not reproducible; reversed in treated cohort | fraction model all ns (\|ρ\| ≤ 0.24); treated cohort ρ = +0.71 (n = 15) |
| Stress confounding of AP-1 | Substantial; survives masking | stress × AP-1 ρ = 0.88 (full, P = 2.3 × 10<sup>−9</sup>), 0.82 (masked, P = 2.5 × 10<sup>−7</sup>), n = 26; masking shrinks histology log2FC 40–64% |
| Histology TF contrasts | MAF higher in LUSC (survives correction); FOXM1 same direction, not significant | MAF P = 0.0013 (adj 0.018; sensitivity-set adj 0.071, provisional); FOXM1 P = 0.006 (adj 0.083) |
| Candidate axes | HGF–MET, FN1–ITGA9, TGFB1 axis (nominal only) | TAM-side P = 0.013, 0.015, 0.058; none survives Bonferroni (adj 0.18, 0.21, 0.81); single cohort, hypothesis-generating |

## Discussion

Three findings qualify the FN1–AP-1 resistance narrative in human tissue. First, the ligand source is CAFs, not macrophages. This is the most robust result in our analysis: it replicates across two cohorts built on different platforms, appears at both the expression level (FN1-positive fractions) and the communication level (ligand–receptor scores), and agrees with the broad understanding that resident fibroblasts are the dominant FN1 producers in epithelial tissues. The macrophage-source model of Ref. [1] is not contradicted at the level of protein localization or mouse genetics, but the human correlative support it rests on is quantitatively consistent with a simpler reading in which macrophage FN1 signal is a minority component of a stroma-dominated program.

Second, the association between FN1 and immune exclusion, which is replicable in bulk data, carries no FN1-specific information. Three independently constructed FN1 signatures, including one that shares no genes with the collagen covariate, lost all association with cytotoxic signatures after ECM adjustment, and neither fraction-nor density-based single-cell measures supported a cellular mechanism. The most parsimonious reading is that FN1 marks collagen-high stromal content, and that stroma-rich tumors are immune-cold for reasons that have little to do with fibronectin as a signaling ligand. This directly limits the clinical extrapolation of murine macrophage-Fn1 knockout studies: the human correlative association they rest on is one that a collagen adjustment eliminates, as shown above.

Third, AP-1 activity, the proposed downstream effector, is substantially confounded by dissociation stress, and the confounding survives the correction we originally set out to test. The near-perfect correlation between sample stress scores and epithelial AP-1 regulon activity (ρ = 0.88) implies that a large share of between-sample AP-1 variance is technical. IEG masking, the correction that would seem most natural, shrank histology-associated effect sizes by 40–64% but left the stress coupling essentially intact (ρ = 0.82), because stress-induced AP-1 activation itself is reflected in non-IEG targets. A third version that also removed the stress genes from the ranking gene universe gave the same value, which rules out a scoring-window artifact and locates the residual signal in the co-expression structure of the regulons rather than in the mechanics of the metric. Because IEG induction is fast, generic and compartment-dependent (strongest in myeloid cells), any single-cell study using AP-1 family members as outcome measures, including studies of immunotherapy resistance, should report IEG-masked versions alongside full-regulon results and, importantly, should not treat masking as sufficient. Reporting a per-sample dissociation-stress score and demonstrating that the headline AP-1 contrast survives adjustment for it is, in our view, the minimum standard for this class of claim.

Several limitations should be acknowledged. GSE148071 contains very few immune cells, so the immune-exclusion question could only be tested in bulk and in the treated cohort; the reverse direction observed there (+0.71) is based on 15 samples with heterogeneous malignant-cell yield and should be read as an absence of support rather than as evidence of a positive FN1–cytotoxicity link. The treated cohort is dominated by post-treatment resections, and response groups are small (four major pathological responders versus eight non-responders among the 12 resected specimens), which limits power for response-level contrasts; FN1-positive fractions did not differ between responders and non-responders in either data type (P = 0.93 for single-cell fractions; P = 0.31 for bulk FN1), consistent with our overall conclusion. Three CNV-positive samples contributed fewer than 50 malignant epithelial cells (11 to 44) and were included in the primary regulon analysis; a prespecified sensitivity analysis restricted to the 23 higher-yield samples left the stress-coupling estimates unchanged but weakened the MAF histology contrast (adjusted P = 0.071). CAF-compartment ligand–receptor scores were available for 17 of 42 GSE148071 samples, so the six-fold communication contrast, while highly significant, rests on the subset of samples with adequate fibroblast capture. Malignant-cell calling was also sensitive to the reference pool used to define the copy-number null. In our implementation the single pathological-complete-responder sample showed no CNV evidence (two affected chromosomes against an immune-cell reference), supporting the built-in specificity check; an independent reimplementation with a different reference pool classified the same sample as CNV-positive. Per-sample malignant-cell counts are likewise implementation-dependent: the 50-cell yield threshold retained 23 of 26 CNV-positive GSE148071 samples in our analysis but 25 in the independent reimplementation. Statistics conditioned on these sets, including the prespecified sensitivity analysis above, should be read with that uncertainty. Sample-set boundaries also affect the candidate-axis contrasts: an independent reimplementation of the full pipeline, using a different malignant-cell pool (40,561 cells across 25 samples versus 43,324 cells across 26 in ours), found TGFB1–TGFBR1/TGFBR2 crossing the Bonferroni threshold in its primary analysis (adjusted P = 0.040, versus 0.81 in ours), so this near-nominal signal in our data should be treated as unstable across plausible analysis sets, and the candidate axes remain hypothesis-generating. Our regulatory-network and communication pipelines are reimplementations rather than the canonical tools, chosen deliberately for transparency and to control the IEG-masking intervention; benchmarking against pySCENIC and established cell–cell communication frameworks on the same inputs would strengthen reproducibility claims. Finally, no spatial or protein-level validation was performed, and the candidate HGF–MET and FN1–ITGA9 contrasts require independent-cohort replication before any therapeutic interpretation.

## Conclusions

In human NSCLC, FN1 is produced predominantly by cancer-associated fibroblasts, and its association with immune exclusion is fully explained by the collagen program of which it forms part; FN1 should be regarded as a marker of collagen-high CAF content rather than an independent therapeutic target. AP-1 measurements in single-cell tumor data are dominated by dissociation-induced immediate-early gene activity; IEG masking is a necessary but insufficient correction, and per-sample stress scores should accompany any AP-1-based resistance claim. HGF–MET and FN1–ITGA9 interactions identified here remain hypothesis-generating candidates for functional study.

## List of abbreviations

AUCell: area under the recovery curve
CAF: cancer-associated fibroblast
CNV: copy-number variation
ECM: extracellular matrix
FN1: fibronectin 1
GEO: Gene Expression Omnibus
IEG: immediate-early gene
LUAD: lung adenocarcinoma
LUSC: lung squamous cell carcinoma
NOS: not otherwise specified
NSCLC: non-small cell lung cancer
PD-1: programmed cell death protein 1
TAM: tumor-associated macrophage
TF: transcription factor
UMI: unique molecular identifier

## Declarations

## Ethics approval and consent to participate

Not applicable. All data were derived from previously published, publicly available de-identified datasets (GEO accessions GSE148071, GSE207422 and GSE131907); the original studies obtained ethics approval and informed consent as reported in their primary publications [6,7,11].

## Consent for publication

Not applicable.

## Data Availability

All datasets analyzed are publicly available at GEO under accessions GSE148071 (https://www.ncbi.nlm.nih.gov/geo/query/acc.cgi?acc=GSE148071) and GSE207422 (https://www.ncbi.nlm.nih.gov/geo/query/acc.cgi?acc=GSE207422) and GSE131907 (https://www.ncbi.nlm.nih.gov/geo/query/acc.cgi?acc=GSE131907). Analysis scripts (Python) and the per-sample result tables supporting the conclusions, including the compartment marker panel used for cell classification, are archived at Zenodo under DOI 10.5281/zenodo.24013282 (https://doi.org/10.5281/zenodo.24013282); the version used for this submission is v1.0.2 (DOI 10.5281/zenodo.24036167).

## Author contributions

ZG: Conceptualization, Data curation, Formal analysis, Funding acquisition, Investigation, Methodology, Project administration, Resources, Software, Supervision, Validation, Visualization, Writing – original draft, Writing – review & editing. WB: Conceptualization, Writing – review & editing. XC: Validation, Writing – review & editing. HJ: Investigation, Writing – review & editing. LX: Data curation, Writing – review & editing. All authors read and approved the final manuscript.

## Acknowledgements

Not applicable.

## Supporting information

**S1 Fig.**
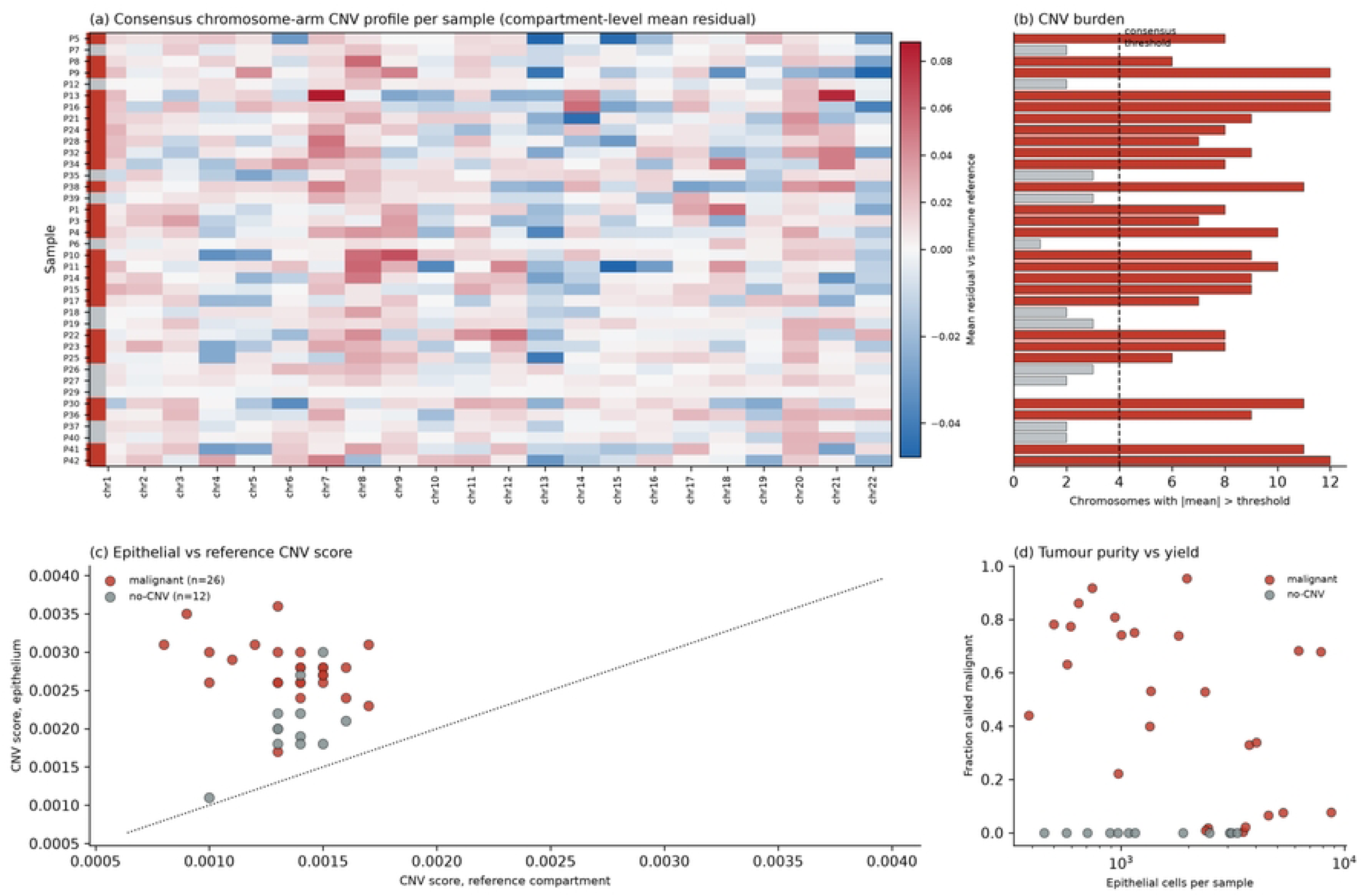
Compartment-level inferCNV consensus: chromosome-arm copy-number profiles for all evaluable samples.

**S2 Fig.**
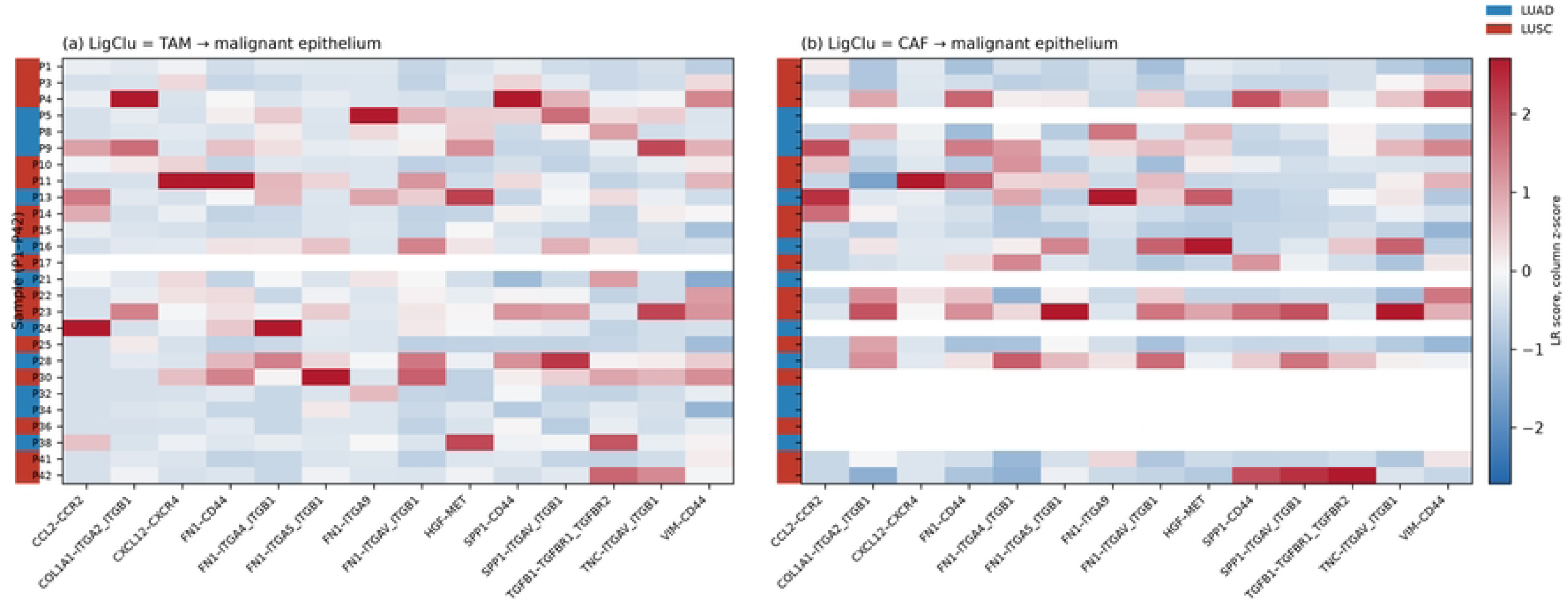
Sample-level ligand-receptor communication scores for the 14 curated axes, by histology.

**S3 Fig.**
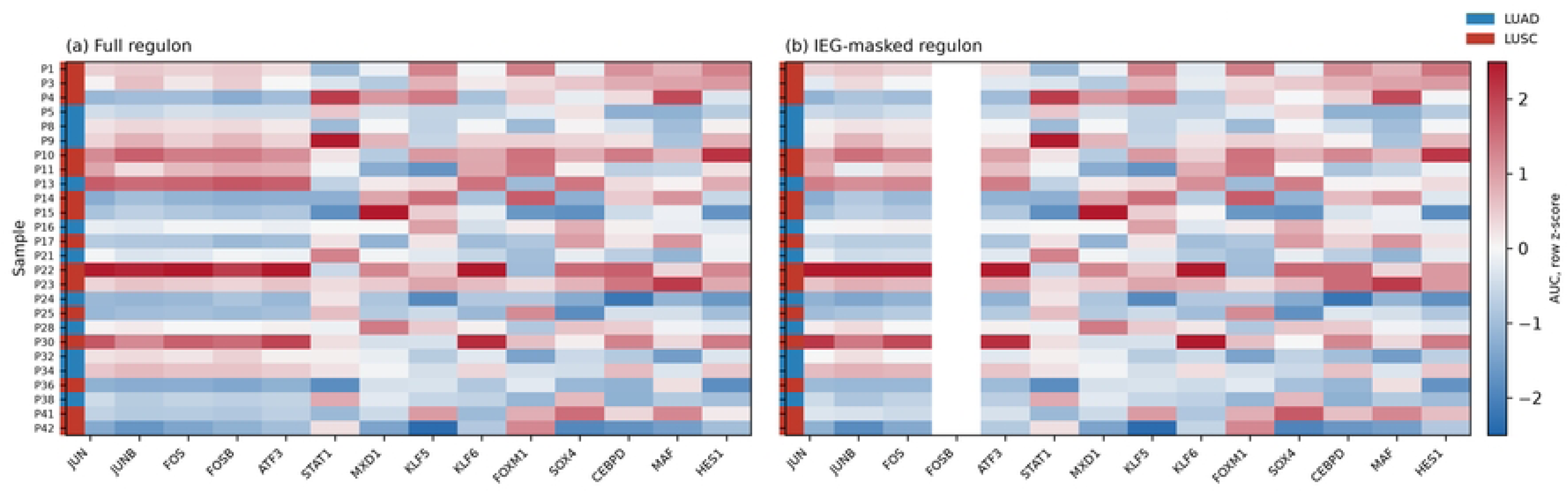
Per-sample regulon activity before and after immediate-early-gene masking.

**S1 Table.** GSE148071 per-sample summary: quality control, compartment composition and malignant-cell yield.

**S2 Table.** GSE207422 per-sample summary with clinical and pathological-response annotations.

**S3 Table.** Regulon statistics for the 14 regulons across the full, masked and hard-masked variants.

**S4 Table.** Ligand-receptor statistics for all candidate axes, by histology.

**S5 Table.** Attribution statistics: raw and partial correlations under each adjustment scheme.

**S6 Table.** Per-sample regulon AUC matrix (full / masked / hard-masked).

**S7 Table.** Per-sample ligand-receptor scores.

**S8 Table.** Hard-mask variant comparison and AUCell window occupancy.

**S9 Table.** GSE131907 third-cohort replication: per-sample FN1-positive fractions and paired Wilcoxon test.

